# GAL4/GFP enhancer-trap lines for identification and manipulation of cells and tissues in developing Arabidopsis leaves

**DOI:** 10.1101/801357

**Authors:** Brindhi Amalraj, Priyanka Govindaraju, Anmol Krishna, Dhruv Lavania, Nguyen Manh Linh, Sree Janani Ravichandran, Enrico Scarpella

## Abstract

Understanding developmental processes requires the unambiguous identification of cells and tissues, and the selective manipulation of the properties of those cells and tissues. Both requirements can most efficiently be satisfied through the use of GAL4/GFP enhancer-trap lines. No such lines are available, however, for the study of leaf development in the Columbia-0 reference genotype of Arabidopsis. Here we address this limitation by identifying and characterizing a set of GAL4/GFP enhancer-trap lines in the Columbia-0 background for the specific labeling of cells and tissues during leaf development, and for the targeted expression of genes of interest in those cells and tissues. By using one line in our set to resolve outstanding problems in leaf vein patterning, we show that these lines can be used to address key questions in plant developmental biology.

## Introduction

Understanding developmental processes requires the unambiguous identification of cells and tissues, and the selective manipulation of the properties of those cells and tissues; both requirements can most efficiently be satisfied by the GAL4 system (Brand & Perrimon, 1993). In this system, a minimal promoter in a construct randomly inserted in a genome responds to neighboring regulatory elements and activates the expression of a gene, included in the same construct, encoding a variant of the GAL4 transcription factor of yeast; the same construct also includes a GAL4-responsive, UAS-driven lacZ, GUS, or GFP, which reports GAL4 expression. Independent, WT-looking lines, in which the construct is inserted in different genomic locations, are selected because they reproducibly express the GAL4-responsive reporter in cell- or tissue-specific patterns. These lines are used to identify cells or tissues, and to drive GAL4-responsive cell- or tissue-specific expression in WT or, through crosses, in mutants and transgenics (e.g., Halder, Callaerts, & Gehring, 1995; Ito, Awano, Suzuki, Hiromi, & Yamamoto, 1997).

The first implementation of the GAL4 system in Arabidopsis was the Haseloff collection of GAL4/GFP enhancer-trap lines, in which an endoplasmic-reticulum-localized GFP (erGFP) responds to the activity of a fusion between the GAL4 DNA-binding domain and the activating domain of the Viral Protein 16 of *Herpex simplex* (Haseloff, 1999). The Haseloff collection is the most extensively used GAL4 system in Arabidopsis (e.g., Sabatini et al., 1999; Sawchuk, Head, Donner, & Scarpella, 2007; Gardner et al., 2009; Wenzel, Marrison, Mattsson, Haseloff, & Bougourd, 2012; Weijers, Van Hamburg, Van Rijn, Hooykaas, & Offringa, 2003; Laplaze et al., 2005), even though it is in the C24 background. This is problematic because the phenotype of hybrids between C24 and Columbia-0 (Col-0), generally considered the reference genotype in Arabidopsis (Koornneef & Meinke, 2010), is different from that of either parent (e.g., Groszmann et al., 2014; Kawanabe et al., 2016; Zhang et al., 2016). The use of GAL4/GFP enhancer-trap lines in the C24 background to investigate processes in the Col-0 background thus imposes the burden of laborious generation of ad-hoc control backgrounds. Therefore, most desirable is the generation and characterization of GAL4/GFP enhancer-trap collections in the Col-0 background. Two such collections have been reported: the Berleth collection, which has been used to identify lines that express GAL4/GFP in vascular tissues (Ckurshumova et al., 2009); and the Poethig collection, which has been used to identify lines that express GAL4/GFP in stomata (Gardner et al., 2009).

Here we screened the Poethig collection and provide a set of lines for the specific labeling of cells and tissues during leaf development, and we show that these lines can be used to address key questions in plant developmental biology.

## Results and Discussion

To identify enhancer-trap lines in the Col-0 background of Arabidopsis with reproducible GAL4-driven GFP expression in developing leaves, we screened the collection generated and donated by Scott Poethig to the Arabidopsis Biological Resource Center. We screened 312 lines for GFP expression in developing leaves; 29 lines satisfied this criterion (Table S1). In 10 of these 29 lines, GFP was expressed in specific cells or tissues; nine of these 10 lines grew normally (Table S1). We imaged GFP expression in first leaves of these nine lines from 2 to 5 days after germination (DAG).

The development of Arabidopsis leaves has been described previously (Pyke, Marrison, & Leech, 1991; Telfer & Poethig, 1994; Kinsman & Pyke, 1998; Candela, Martinez-Laborda, & Micol, 1999; Donnelly, Bonetta, Tsukaya, Dengler, & Dengler, 1999; Mattsson, Sung, & Berleth, 1999; Kang & Dengler, 2002; Kang & Dengler, 2004; Mattsson, Ckurshumova, & Berleth, 2003; Scarpella, Francis, & Berleth, 2004; Larkin, Young, Prigge, & Marks, 1996). Briefly, at 2 DAG the first leaf is recognizable as a cylindrical primordium with a midvein at its center (Fig. 1A). By 2.5 DAG, the primordium has elongated and expanded (Fig. 1B). By 3 DAG, the primordium has continued to expand, and the first loops of veins (“first loops”) have formed (Fig. 1C). By 4 DAG, a lamina and a petiole have become recognizable, second loops have formed, and minor veins have started to form in the top half of the lamina (Fig. 1D). By 5 DAG, lateral outgrowths (hydathodes) have become recognizable in the bottom quarter of the lamina, third loops have formed, and minor vein formation has spread toward the base of the lamina (Fig. 1E). Leaf hairs (trichomes) and pores (stomata) can be first recognized at the tip of 2.5-and 3-DAG primordia, respectively, and their formation spreads toward the base of the lamina during leaf development (Fig. 1F–I).

**Figure 1.**
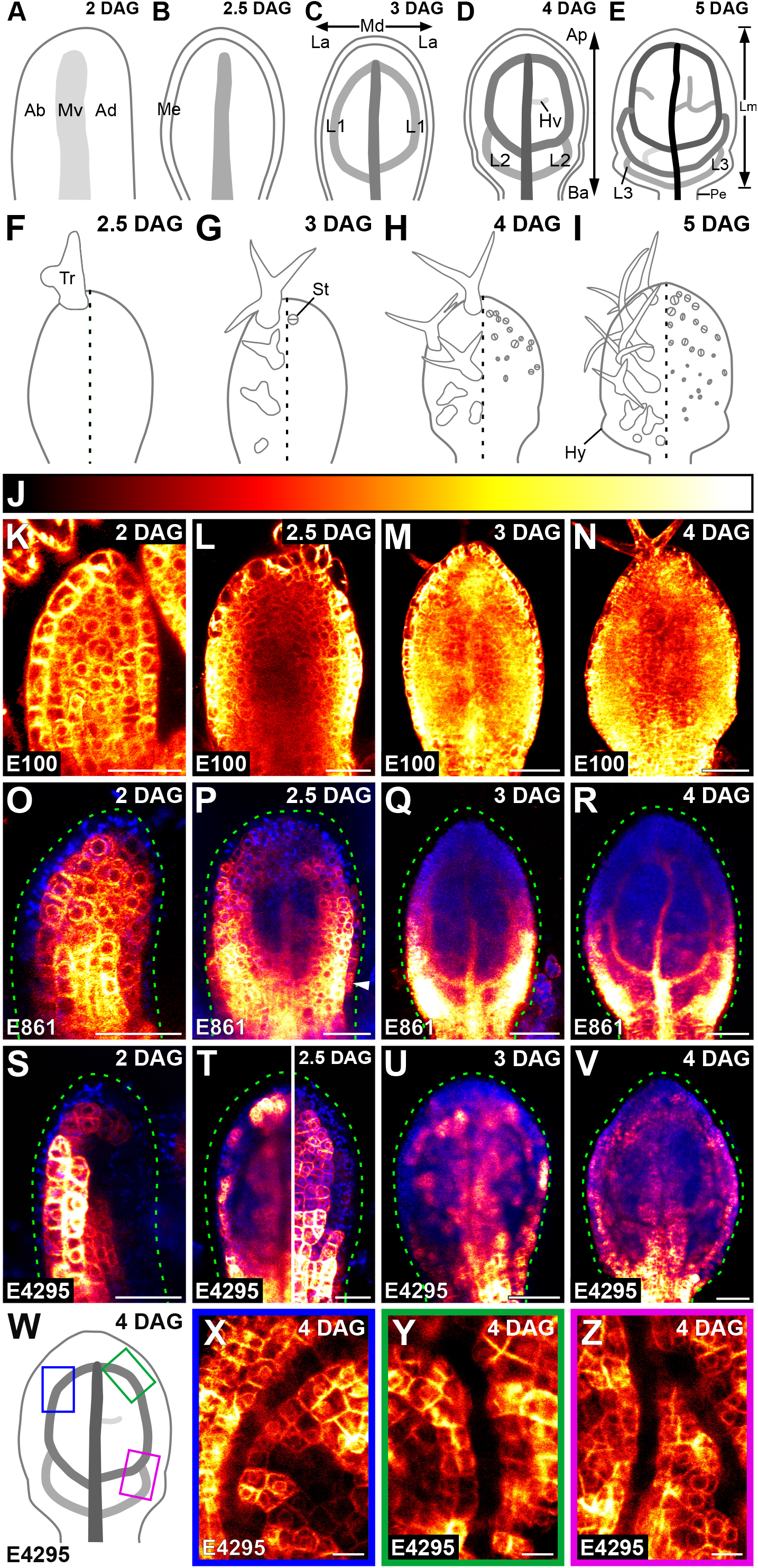
Expression of E100>>, E861>> and E4295>>erGFP in Arabidopsis Leaf Development. (A–Z) First leaves. Top right: leaf age in days after germination (DAG). (A–E) Development of leaf and veins; increasingly darker grays depict progressively later stages of vein development. See text for details. (F–I) Development of trichomes and stomata in adaxial (left) or abaxial (right) epidermis. See text for details. Ab: abaxial; Ad: adaxial; Ap: apical; Ba: basal; Hv: minor vein; Hy: hydathode; L1, L2 and L3: first, second and third loop; La: lateral; Lm: lamina; Md: median; Me: marginal epidermis; Mv: midvein; Pe: petiole; St: stoma; Tr: trichome. (K–V,X– Z) Confocal laser scanning microscopy. Bottom left: genotype. Look–up table (ramp in J) visualizes erGFP expression levels. Blue: autofluorescence. Dashed green line delineates leaf outline. White arrowhead points to epidermal expression. (K–S,U,V,X–Z) Median view (abaxial side to the left in K). (T) Median (left) and abaxial subepidermal (right) views. (W) Increasingly darker grays depict progressively later stages of vein development. Boxes illustrate positions of closeups in X, Y and Z. See Table S2 for reproducibility of expression features. Bars: (K,L,O,P,S,T) 30 μm; (M,N,Q,R,U,V) 60 μm; (X–Z) 10 μm.

Consistent with previous observations (Huang et al., 2014), E100>>erGFP was expressed in all the cells of 2-, 2.5-, 3-, and 4-DAG leaf primordia (Fig. 1K– N).

Consistent with previous observations (Krogan & Berleth, 2012), E861>>erGFP was expressed in all the inner cells of the 2-DAG primordium, though more strongly in its innermost cells (Fig. 1O). At 2.5 DAG, expression had been activated in the lowermost epidermal cells of the primordium margin and persisted in all the inner cells of the bottom half of the primordium; in the top half of the primordium, weaker expression persisted in inner cells, except near the midvein, where by then it had been terminated (Fig. 1P). At 3 DAG, expression continued to persist in all the inner cells of the bottom half of the primordium, though expression was stronger in the areas where second loops were forming; in the top half of the primordium, weaker expression had become restricted to the midvein, first loops and minor veins (Fig. 1Q). At 4 DAG, expression in the top half of the leaf remained restricted to the midvein, first loops and minor veins, and in the bottom half of the leaf it had declined in inner cells between the first loops and the developing second loops (Fig. 1R). In summary, E861>>erGFP was expressed ubiquitously at early stages of inner-cell development; over time, however, expression became restricted to developing veins. As such, expression of E861>>erGFP closely resembles that of *MONOPTEROS* and *PIN-FORMED1*, which marks the gradual selection of vascular cells from within the leaf inner tissue (Scarpella, Marcos, Friml, & Berleth, 2006; Wenzel, Schuetz, Yu, & Mattsson, 2007).

E4295>>erGFP expression was restricted to inner cells in 2-, 2.5-, 3-, and 4-DAG leaf primordia (Fig. 1S–V,X–Z). At 2 DAG, E4295>>erGFP was expressed almost exclusively in the inner cells of the abaxial side of the primordium (Fig. 1S), but by 2.5 DAG it had spread to the middle tissue layer (Fig. 1T), from which veins form (Stewart, 1978; Tilney-Bassett, 1986). Expression persisted in the inner cells of the abaxial side and of the middle tissue layer in 3-and 4-DAG primordia (Fig. 1U,V). High-resolution images of the middle tissue layer showed that expression was excluded from developing veins (Fig. 1X–Z), suggesting that it marks inner, non-vascular cells. Therefore, expression of E4295>>erGFP closely resembles that of *LIGHT HARVESTING COMPLEX A6* and *SCARECROW-LIKE32* (Sawchuk, Donner, Head, & Scarpella, 2008; Gardiner, Donner, & Scarpella, 2011), and that of J0571>>erGFP in the C24 background (Wenzel et al., 2012).

At 2 DAG, E4259>>erGFP was expressed in the top third of the median adaxial epidermis and in the whole median abaxial epidermis, though expression was stronger in the top half of the primordium (Fig. 2A). By 2.5 DAG, strong expression had spread to the whole abaxial and to the top three-quarters of the marginal epidermis; expression had spread to the top three-quarters of the adaxial epidermis too, but it was stronger in the top half of the primordium (Fig. 2B,F). At 3 DAG, strong expression had spread to the top three-quarters of the adaxial epidermis and to the whole marginal epidermis, and persisted in the whole abaxial epidermis (Fig. 2C,G). At 4 DAG, expression persisted in the whole marginal epidermis, continued to persist in the whole abaxial epidermis, and had spread to the whole lamina and the petiole midline in the adaxial epidermis (Fig. 2D,H). At all analyzed stages, E4259>>erGFP was expressed in trichomes but was not expressed in mature stomata (Fig. 2B–H). In conclusion, expression of E4259>>erGFP closely resembles that of *ARABIDOPSIS THALIANA MERISTEM LAYER1* (Lu, Porat, Nadeau, & O’Neill, 1996; Sessions, Weigel, & Yanofsky, 1999), which marks epidermal cells and whose promoter is used to drive epidermis-specific expression (e.g., Kierzkowski, Lenhard, Smith, & Kuhlemeier, 2013; Bilsborough et al., 2011; Takada & Jürgens, 2007).

**Figure 2.**
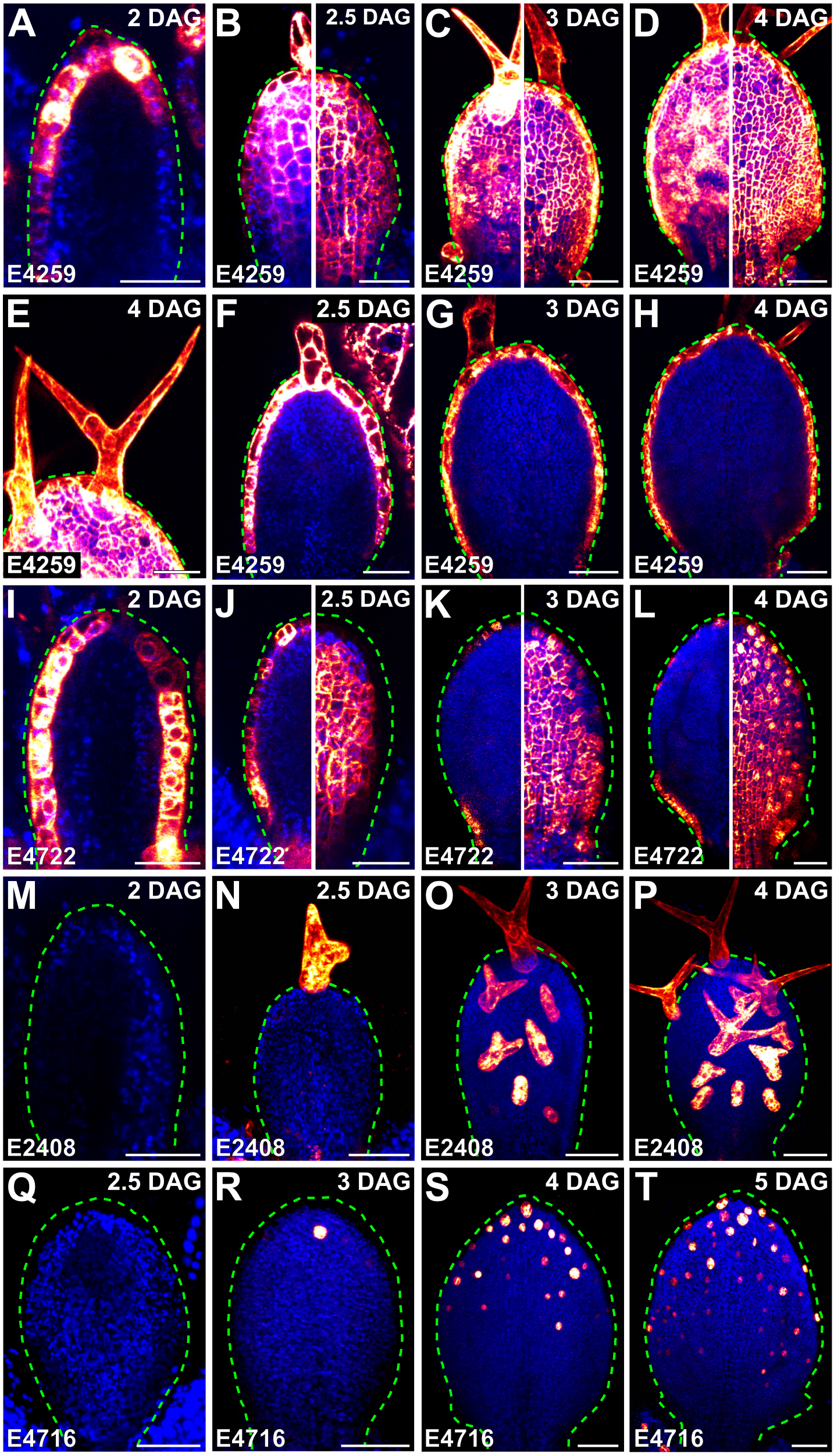
Expression of E4259>>, E4722>>, E2408>> and E4716>>erGFP in Leaf Development. (A–T) Confocal laser scanning microscopy. First leaves. Top right: leaf age in days after germination (DAG). Bottom left: genotype. Look–up table (ramp in Fig. 1J) visualizes erGFP expression levels. Blue: autofluorescence. Dashed green line delineates leaf outline. (A,F–I,M) Median view (abaxial side to the left in A,I,M). (B–D) Adaxial (left) and abaxial (right) epidermal views. (E) Closeup of trichome in D, left. (J–L) Median (left) and abaxial epidermal (right) views. (N–P) Adaxial epidermal view. (Q–T) Abaxial epidermal view. See Table S2 for reproducibility of expression features. Bars: (A,B,F,I,J,M,N,Q) 30 μm; (C,D,E,G,H,K,L,O,P,R,S,T) 60 μm.

E4722>>erGFP was expressed in all the epidermal cells of the 2-DAG primordium, though more weakly at its tip (Fig. 2I). E4722>>erGFP was expressed in all the epidermal cells of the 2.5-DAG primordium too, except at its margin, where expression had been terminated in a few cells of its top half (Fig. 2J). At 3 DAG, expression persisted in all the epidermal cells, except at the primordium margin, where expression had been terminated in most of the cells of its top three-quarters (Fig. 2K). At 4 DAG, expression continued to persist in all the epidermal cells, except at the leaf margin, where expression had almost completely been terminated in the cells of its top three-quarters (Fig. 2L). Unlike E4259>>erGFP, E4722>>erGFP was expressed in stomata but was not expressed in trichomes (Fig. 2J–L).

At all analyzed stages, expression of E2408>>erGFP and E4716>>erGFP was restricted to trichomes and stomata, respectively. E2408>>erGFP was first expressed in developing trichomes at the tip of the 2.5-DAG primordium (Fig. 2M,N). By 3 DAG, expression had spread to developing and mature trichomes in the top three-quarters of the primordium (Fig. 2O), and by 4 DAG to those in the whole lamina (Fig. 2P). E4716>>erGFP was first expressed in stomata at the tip of the 3-DAG primordium (Fig. 2Q,R). By 4 DAG, expression had spread to the stomata in the top half of the lamina (Fig. 2S), and by 5 DAG to those in its top three-quarters (Fig. 2T).

At all analyzed stages, expression of E2331>>erGFP and E3912>>erGFP was restricted to developing veins. E2331>>erGFP was expressed in both isodiametric and elongated cells of the midvein in 2-and 2.5-DAG primordia (Fig. 3A,B). By 3 DAG, it was expressed in first loops, and by 4 DAG in second loops and minor veins (Fig. 3C,D). E3912>>erGFP was first expressed in the midvein of the 3-DAG primordium (Fig. 3E,F). By 4 DAG, expression had spread to first loops, and by 5 DAG to second loops and minor veins (Fig. 3G,H). These observations suggest that expression of E3912>>erGFP is initiated later than that of E2331>>erGFP in vein development. Furthermore, because the expression of E2331>>erGFP appears to be no different from that of the preprocambial markers ATHB8::nYFP, J1721>>erGFP and SHR::nYFP (Sawchuk et al., 2007; Donner, Sherr, & Scarpella, 2009; Gardiner et al., 2011), we suggest that E2331>>erGFP expression marks preprocambial stages of vein development, a conclusion that is consistent with E2331>>erGFP expression during embryogenesis (Gillmor et al., 2010). Finally, because E3912>>erGFP expression appears to be no different from that of the procambial marker Q0990>>erGFP in the C24 background (Sawchuk et al., 2007), we suggest that E3912>>erGFP expression marks procambial stages of vein development.

**Figure 3.**
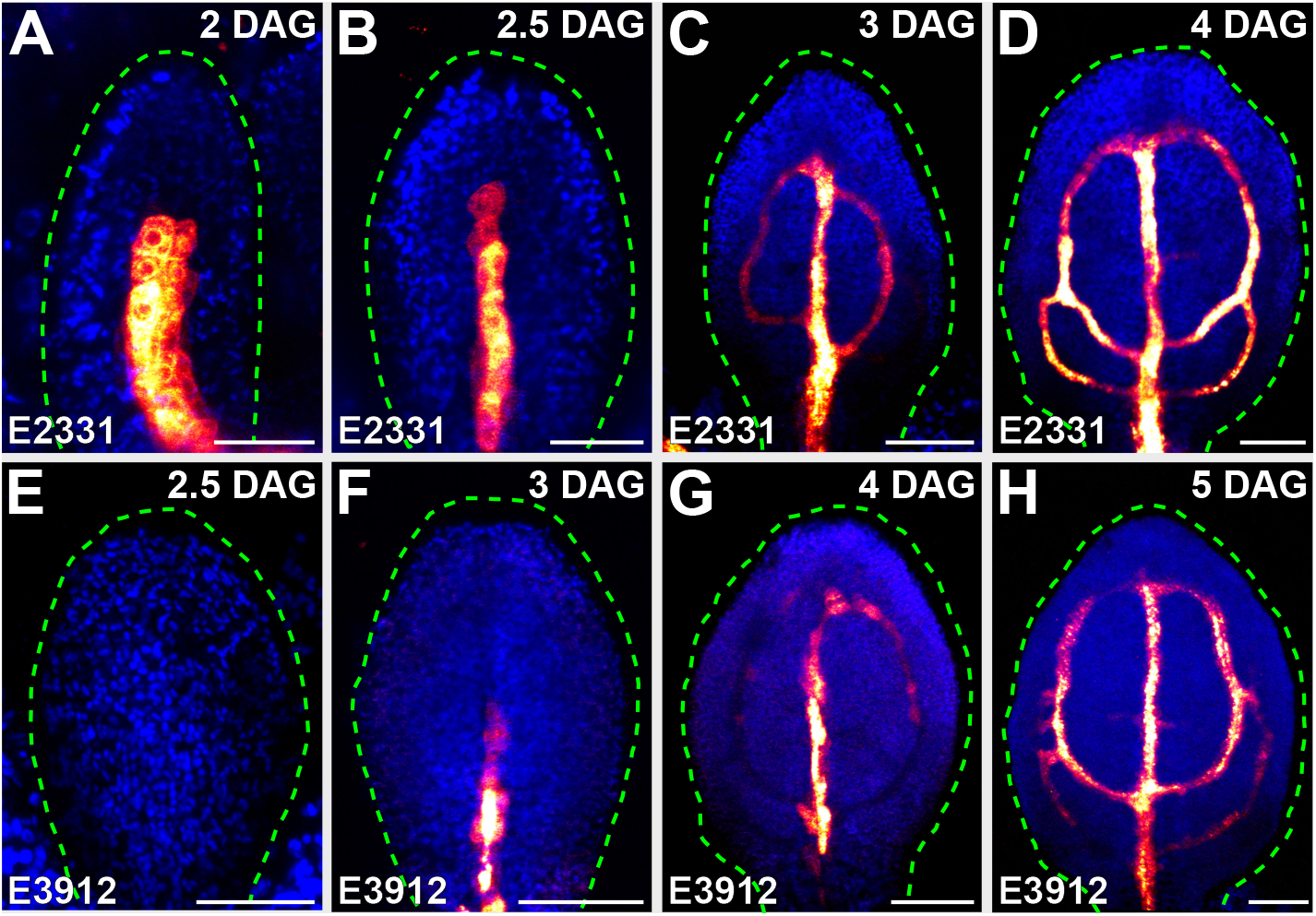
Expression of E2331>> and E3912>>erGFP in Leaf Development. (A–H) Confocal laser scanning microscopy. First leaves. Top right: leaf age in days after germination (DAG). Bottom left: genotype. Look–up table (ramp in Fig. 1J) visualizes erGFP expression levels. Blue: autofluorescence. Dashed green line delineates leaf outline. Median view (abaxial side to the left in A). See Table S2 for reproducibility of expression features. Bars: (A,B,E) 30 μm; (C,D,F–H) 60 μm.

To show the informative power of the lines reported here for plant developmental biology, we selected the E2331 line, which marks early stages of vein development (Fig. 3A–D).

In WT leaves, the elongated vascular cells are connected to one another into continuous veins (Esau, 1965) (Fig. 4A). By contrast, in mature leaves of the *gnom* (*gn*) mutant, putative vascular cells fail to elongate and to connect to one another into continuous veins; instead, they accumulate into shapeless clusters of seemingly disconnected and randomly oriented cells (Shevell, Kunkel, & Chua, 2000; Verna, Ravichandran, Sawchuk, Linh, & Scarpella, 2019) (Fig. 4B). Though the cells in these clusters have some features of vascular cells (e.g., distinctive patterns of secondary cell-wall thickenings), they lack others (e.g., elongated shape and end-to-end connection to form continuous veins). Therefore, it is unclear whether the clustered cells in *gn* mature leaves are abnormal vascular cells or nonvascular cells that have recruited a cellular differentiation pathway that is normally, but not always (e.g., Solereder, 1908; Kubo et al., 2005; Yamaguchi et al., 2010), associated with vascular development. To address this question, we imaged E2331>>erGFP expression in developing leaves of WT and *gn*.

**Figure 4.**
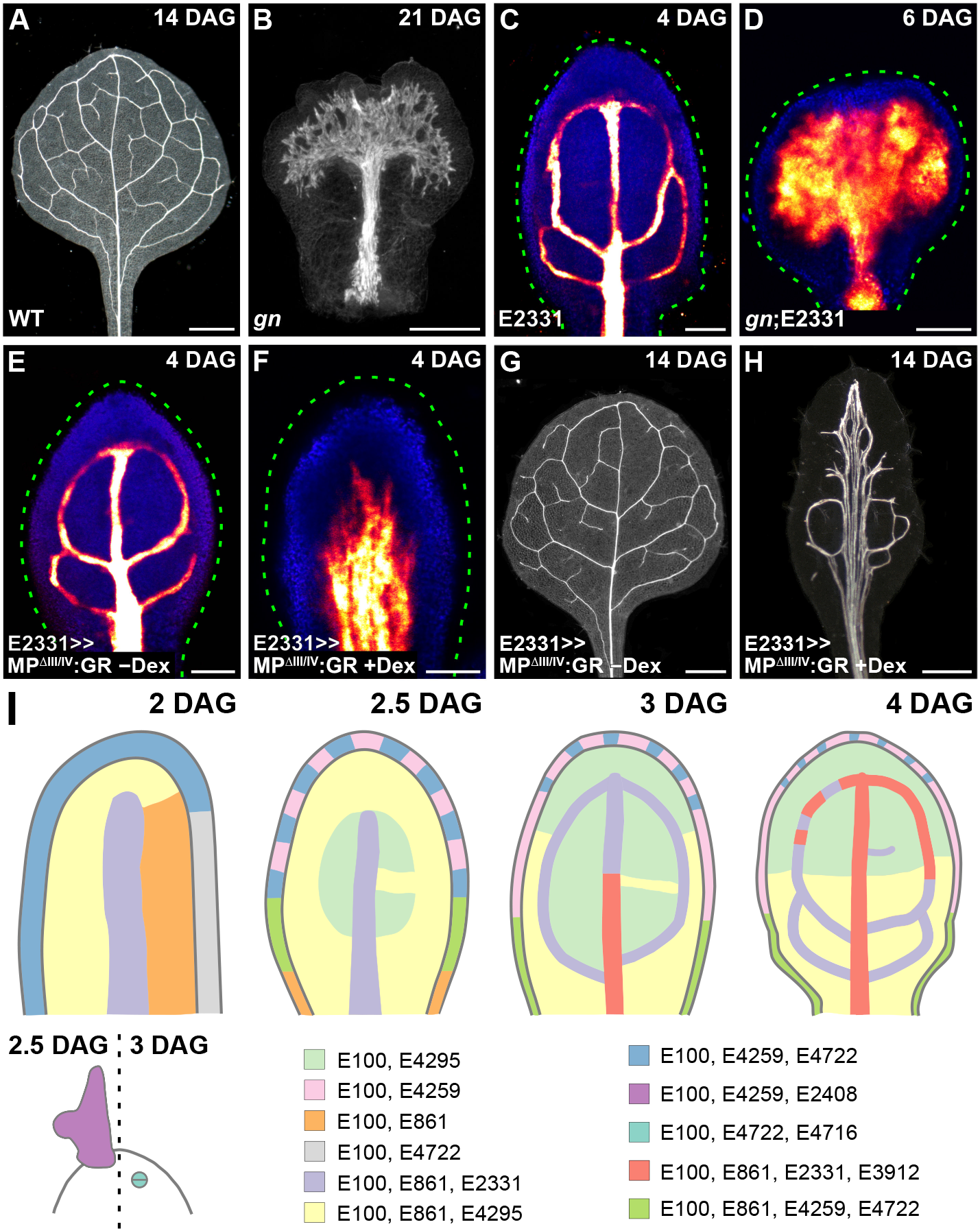
E2331-Mediated Visualization and Manipulation of Developing Veins. (A–H) First leaves. Top right: leaf age in days after germination (DAG). Bottom left: genotype and treatment. (A,B,G,H) Dark-field microscopy of cleared leaves. (C–F) Confocal laser scanning microscopy. Look–up table (ramp in Fig. 1J) visualizes erGFP expression levels. Blue: autofluorescence. Dashed green line delineates leaf outline. Median view. See Table S2 for reproducibility of expression and pattern features. (I) Expression map of E100>>, E861>>, E4295>>, E4259>>, E4722>>, E2408>>, E4716>>, E2331>> and E3912>>erGFP in leaf development. See text for details. Bars: (A,B,G,H) 500 μm; (C–F) 60 μm.

As shown above (Fig. 3D), E2331>>erGFP was expressed in midvein, first and second loops, and minor veins in WT (Fig. 4C). In *gn*, the pattern of E2331>>erGFP expression in developing leaves recapitulated that of vascular differentiation in mature leaves (Fig. 4B,D), suggesting that the putative vascular cells in the shapeless clusters are indeed vascular cells, albeit abnormal ones.

Auxin signaling is thought to be required for vein formation because mutations in genes involved in auxin signaling or treatment with inhibitors of auxin signaling leads to the formation of fewer, incompletely differentiated veins (Przemeck, Mattsson, Hardtke, Sung, & Berleth, 1996; Hardtke & Berleth, 1998; Mattsson et al., 2003; Verna et al., 2019). Increasing auxin signaling by means of broadly expressed mutations or transgenes turns nearly every cell file in the developing leaf into a vein, suggesting that auxin signaling is also sufficient for vein formation (Krogan, Ckurshumova, Marcos, Caragea, & Berleth, 2012; Garrett et al., 2012). This interpretation assumes that it is the increased auxin signaling in the cell files that normally would not differentiate into veins that leads those cell files to differentiate in fact into veins. However, it is also possible that it is the increased auxin signaling in the cell files that normally differentiate into veins that leads the flanking cell files, which normally would not differentiate into veins, to do in fact so. To discriminate between these possibilities, we increased auxin signaling in developing veins by expressing by the E2331 driver a dexamethasone (dex)-inducible MPΔIII/IV (Krogan et al., 2012; Ckurshumova, Smirnova, Marcos, Zayed, & Berleth, 2014; Smetana et al., 2019) (MPΔIII/IV:GR), and we imaged E2331>>erGFP expression in developing leaves and vein patterns in mature leaves of E2331>>MPΔIII/IV:GR grown with or without dex.

Consistent with previous observations (Fig. 3D; Fig. 4C), in developing leaves of E2331>>MPΔIII/IV:GR grown without dex, E2331>>erGFP was expressed in narrow domains (Fig. 4E). By contrast, E2331>>erGFP was expressed in broad domains in developing leaves of dex-grown E2331>>MPΔIII/IV:GR (Fig. 4F). Whether with or without dex, the patterns of E2331>>erGFP expression in developing leaves of E2331>>MPΔIII/IV:GR presaged those of vein formation in mature leaves: narrow zones of vein formation in the absence of dex; broad areas of vascular differentiation in the presence of dex, often with multiple veins running parallel next to one another (Fig. 4G,H). Though the areas of vascular differentiation in dex-grown E2331>>MPΔIII/IV:GR are not as broad as those of leaves in which MPΔIII/IV is expressed in all the inner cells (Krogan et al., 2012), they are broader than those of E2331>>MPΔIII/IV:GR grown without dex. These observations suggest that, at least in part, it is the increased auxin signaling in the cell files that normally differentiate into veins that leads the flanking cell files, which normally would not differentiate into veins, to do in fact so.

In conclusion, we provide a set of GAL4/GFP enhancer-trap lines in the Col-0 background of Arabidopsis for the specific labeling of cells and tissues during leaf development (Fig. 4I), and we show that these lines can be used to address key questions in plant developmental biology.

## Materials & Methods

### Plants

Origin and nature of GAL4 enhancer-trap lines are in Table S1. *gn-13* (SALK_045424; ABRC) (Alonso et al., 2003) (Verna et al., 2019) contains a T-DNA insertion after nucleotide +2835 of *GN* and was genotyped with the “SALK_045424 gn LP” (5’-TGATCCAAATCACTGGGTTTC-3’) and “SALK_045424 gn RP” (5’-AGCTGAAGATAGGGAATTCGC-3’) oligonucleotides (*GN*) and with the “SALK_045424 gn RP” and “LBb1.3” (5’-ATTTTGCCGATTTCGGAAC-3’) oligonucleotides (*gn*). To generate the UAS::MPΔIII/IV:GR construct, the UAS promoter was amplified with the “UAS Promoter SalI Forward” (5’-ATAGTCGACCCAAGCGCGCAATTAACCCTCAC-3’) and the “UAS Promoter XhoI Reverse” (5’-AGCCTCGAGCCTCTCCAAATGAAATGAACTTCC-3’) oligonucleotides; MPΔIII/IV was amplified with the “MP Delta XhoI Forward” (5’-AAACTCGAGATGATGGCTTCATTGTCTTGTGTT-3’) and the “MP EcoRI Reverse” (5’-ATTGAATTCGGTTCGGACGCGGGGTGTCGCAATT-3’) oligonucleotides; and a fragment of the rat glucocorticoid (GR) receptor gene was amplified with the “SpeI GR Forward” (5-‘GGGACTAGTGGAGAAGCTCGAAAAACAAAG-3’) and the “GR ApaI Reverse” (5’-GCGGGGCCCTCATTTTTGATGAAACAG-3’) oligonucleotides. Seeds were sterilized and sown as in (Sawchuk et al., 2008). Stratified seeds were germinated and seedlings were grown at 22°C under continuous fluorescent light (~80 μmol m^−2^ s^−1^). Plants were grown at 24°C under fluorescent light (~85 μmol m^−2^ s^−1^) in a 16–h–light/8–h–dark cycle. Plants were transformed and representative lines were selected as in (Sawchuk et al., 2008).

### Chemicals

Dexamethasone (Sigma-Aldrich, catalogue no. D4902) was dissolved in dimethyl sulfoxide and was added to growth medium just before sowing.

### Imaging

Developing leaves were mounted and imaged as in (Sawchuk, Edgar, & Scarpella, 2013), except that emission was collected from ~1.5–5-μm-thick optical slices. Fluorophores were excited with the 488-nm line of a 30-mW Ar laser; GFP emission was collected with a BP 505–530 filter, and autofluorescence was collected between 550 and 754 nm. Mature leaves were fixed in 3 : 1 or 6 : 1 ethanol : acetic acid, rehydrated in 70% ethanol and in water, cleared briefly (few seconds to few minutes) — when necessary — in 0.4 M sodium hydroxide, washed in water, mounted in 80% glycerol or in 1 : 2 : 8 or 1 : 3 : 8 water : glycerol : chloral hydrate, and imaged as in (Odat et al., 2014). In the Fiji distribution (Schindelin et al., 2012) of ImageJ (Schindelin, Rueden, Hiner, & Eliceiri, 2015; Rueden et al., 2017; Schneider, Rasband, & Eliceiri, 2012), grayscaled RGB color images were turned into 8-bit images; when necessary, 8-bit images were combined into stacks, and maximum-intensity projection was applied to stacks; look-up-tables were applied to images or stacks, and brightness and contrast were adjusted by linear stretching of the histogram.

## Acknowledgements

We thank the Arabidopsis Biological Resource Center for seeds. This work was supported by Discovery Grants of the Natural Sciences and Engineering Research Council of Canada (NSERC) to E.S. A.K. was supported by an NSERC CGS-M Scholarship.

