## Supplemental Table 1 for "GAL4/GFP enhancer-trap lines for identification and manipulation of cells and tissues in developing Arabidopsis leaves"

**Table S1. Origin and nature of lines**

| **ABRC stock no.** | **Donor stock no.** | **Expression in developing leaves** | **Tissue- and/or stage-specific expression** | **Wild-type looking** |
| --- | --- | --- | --- | --- |
| CS24240 | E53 | N | ··· | ··· |
| CS24241 | E306 | N | ··· | ··· |
| CS24242 | E337 | N | ··· | ··· |
| CS24243 | E362 | N | ··· | ··· |
| CS24244 | E456 | N | ··· | ··· |
| CS24245 | E513 | N | ··· | ··· |
| CS24246 | E652 | N | ··· | ··· |
| CS24247 | E751 | N | ··· | ··· |
| CS24248 | E788 | N | ··· | ··· |
| CS24249 | E829 | N | ··· | ··· |
| CS24250 | E1012 | N | ··· | ··· |
| CS24251 | E1075 | N | ··· | ··· |
| CS24252 | E1195 | N | ··· | ··· |
| CS24253 | E1247 | N | ··· | ··· |
| CS24254 | E1287 | N | ··· | ··· |
| CS24255 | E1324 | N | ··· | ··· |
| CS24256 | E1332 | Y | N | ··· |
| CS24257 | E2042 | N | ··· | ··· |
| CS24258 | E2065 | N | ··· | ··· |
| CS24259 | E2072 | N | ··· | ··· |
| CS24260 | E2119 | N | ··· | ··· |
| CS24262 | E2168 | N | ··· | ··· |
| CS24264 | E2242 | N | ··· | ··· |
| CS24265 | E2263 | N | ··· | ··· |
| CS24266 | E2271 | N | ··· | ··· |
| CS24267 | E2306 | N | ··· | ··· |
| CS24269 | E3191 | N | ··· | ··· |
| CS24270 | E3597 | N | ··· | ··· |
| CS24271 | E3604 | N | ··· | ··· |
| CS24272 | E4259 | Y | Y | Y |
| CS65892 | E2331 | Y | Y | Y |
| CS65893 | E2023 | N | ··· | ··· |
| CS67882 | suo-1 | N | ··· | ··· |
| CS70001 | E1 | N | ··· | ··· |
| CS70002 | E3 | N | ··· | ··· |
| CS70003 | E63 | N | ··· | ··· |
| CS70004 | E66 | N | ··· | ··· |
| CS70005 | E74 | Y | N | ··· |
| CS70006 | E829 | N | ··· | ··· |
| CS70007 | E100 | Y | Y | Y |
| CS70008 | E103 | N | ··· | ··· |
| CS70009 | E105 | N | ··· | ··· |
| CS70010 | E107 | N | ··· | ··· |
| CS70011 | E135 | N | ··· | ··· |
| CS70012 | E144 | N | ··· | ··· |
| CS70013 | E183 | N | ··· | ··· |
| CS70014 | E191 | N | ··· | ··· |
| CS70015 | E226 | N | ··· | ··· |
| CS70016 | E227 | Y | N | ··· |
| CS70017 | E230 | N | ··· | ··· |
| CS70018 | E232 | N | ··· | ··· |
| CS70019 | E242 | N | ··· | ··· |
| CS70020 | E244 | N | ··· | ··· |
| CS70021 | E254 | N | ··· | ··· |
| CS70022 | E259 | Y | N | ··· |
| CS70023 | E268 | N | ··· | ··· |
| CS70024 | E280 | N | ··· | ··· |
| CS70025 | E292 | N | ··· | ··· |
| CS70026 | E314 | N | ··· | ··· |
| CS70027 | E325 | N | ··· | ··· |
| CS70028 | E336 | N | ··· | ··· |
| CS70029 | E340 | Y | N | ··· |
| CS70030 | E361 | N | ··· | ··· |
| CS70031 | E387 | N | ··· | ··· |
| CS70032 | E434 | N | ··· | ··· |
| CS70033 | E457 | N | ··· | ··· |
| CS70034 | E461 | N | ··· | ··· |
| CS70035 | E462 | N | ··· | ··· |
| CS70036 | E464 | N | ··· | ··· |
| CS70037 | E470 | N | ··· | ··· |
| CS70038 | E491 | N | ··· | ··· |
| CS70039 | E555-1 | N | ··· | ··· |
| CS70040 | E555-2 | N | ··· | ··· |
| CS70041 | E556 | N | ··· | ··· |
| CS70042 | E583 | N | ··· | ··· |
| CS70043 | E655 | N | ··· | ··· |
| CS70044 | E657 | Y | N | ··· |
| CS70045 | E658 | N | ··· | ··· |
| CS70046 | E668 | N | ··· | ··· |
| CS70047 | E698 | N | ··· | ··· |
| CS70048 | E700 | N | ··· | ··· |
| CS70049 | E719 | N | ··· | ··· |
| CS70050 | E744 | N | ··· | ··· |
| CS70051 | E771 | N | ··· | ··· |
| CS70052 | E790 | N | ··· | ··· |
| CS70053 | E835 | N | ··· | ··· |
| CS70054 | E838 | N | ··· | ··· |
| CS70055 | E861 | Y | Y | Y |
| CS70056 | E864 | N | ··· | ··· |
| CS70057 | E876 | N | ··· | ··· |
| CS70058 | E884 | N | ··· | ··· |
| CS70059 | E892 | N | ··· | ··· |
| CS70060 | E894 | N | ··· | ··· |
| CS70061 | E903 | N | ··· | ··· |
| CS70062 | E910 | N | ··· | ··· |
| CS70063 | E912 | N | ··· | ··· |
| CS70065 | E939 | N | ··· | ··· |
| CS70066 | E940 | N | ··· | ··· |
| CS70067 | E945 | N | ··· | ··· |
| CS70068 | E951 | N | ··· | ··· |
| CS70069 | E992 | N | ··· | ··· |
| CS70070 | E994 | N | ··· | ··· |
| CS70071 | E1049 | N | ··· | ··· |
| CS70072 | E1092 | N | ··· | ··· |
| CS70073 | E1100 | N | ··· | ··· |
| CS70074 | E1127 | N | ··· | ··· |
| CS70075 | E1128 | N | ··· | ··· |
| CS70076 | E1130 | N | ··· | ··· |
| CS70077 | E1155 | N | ··· | ··· |
| CS70078 | E1161 | N | ··· | ··· |
| CS70079 | E1176 | N | ··· | ··· |
| CS70080 | E1222 | N | ··· | ··· |
| CS70081 | E1223 | N | ··· | ··· |
| CS70082 | E1237 | N | ··· | ··· |
| CS70083 | E1238 | N | ··· | ··· |
| CS70084 | E1250 | N | ··· | ··· |
| CS70085 | E1252 | N | ··· | ··· |
| CS70086 | E1271 | N | ··· | ··· |
| CS70087 | E1289 | Y | N | ··· |
| CS70088 | E1304 | N | ··· | ··· |
| CS70089 | E1322 | N | ··· | ··· |
| CS70090 | E1325 | N | ··· | ··· |
| CS70091 | E1331 | N | ··· | ··· |
| CS70092 | E1341 | N | ··· | ··· |
| CS70093 | E1344 | N | ··· | ··· |
| CS70094 | E1356 | N | ··· | ··· |
| CS70095 | E1361 | N | ··· | ··· |
| CS70096 | E1362 | N | ··· | ··· |
| CS70097 | E1370 | N | ··· | ··· |
| CS70098 | E1387 | N | ··· | ··· |
| CS70099 | E1388 | N | ··· | ··· |
| CS70100 | E1395 | N | ··· | ··· |
| CS70101 | E1396 | N | ··· | ··· |
| CS70102 | E1405 | N | ··· | ··· |
| CS70103 | E1416 | N | ··· | ··· |
| CS70104 | E1439 | N | ··· | ··· |
| CS70105 | E1439m | N | ··· | ··· |
| CS70106 | E1457 | N | ··· | ··· |
| CS70107 | E1567 | N | ··· | ··· |
| CS70108 | E1570 | N | ··· | ··· |
| CS70109 | E1607 | N | ··· | ··· |
| CS70110 | E1626 | N | ··· | ··· |
| CS70111 | E1627 | N | ··· | ··· |
| CS70112 | E1628 | N | ··· | ··· |
| CS70113 | E1638 | N | ··· | ··· |
| CS70114 | E1644 | N | ··· | ··· |
| CS70115 | E1662 | N | ··· | ··· |
| CS70116 | E1663 | Y | N | ··· |
| CS70117 | E1665 | N | ··· | ··· |
| CS70118 | E1678 | N | ··· | ··· |
| CS70119 | E1684 | N | ··· | ··· |
| CS70120 | E1689 | N | ··· | ··· |
| CS70121 | E1691 | N | ··· | ··· |
| CS70122 | E1701 | N | ··· | ··· |
| CS70123 | E1728 | N | ··· | ··· |
| CS70125 | E1751 | N | ··· | ··· |
| CS70126 | E1765 | N | ··· | ··· |
| CS70127 | E1767 | N | ··· | ··· |
| CS70128 | E1785 | N | ··· | ··· |
| CS70129 | E1786 | N | ··· | ··· |
| CS70130 | E1797 | N | ··· | ··· |
| CS70131 | E1801 | N | ··· | ··· |
| CS70132 | E1809 | N | ··· | ··· |
| CS70133 | E1815 | N | ··· | ··· |
| CS70134 | E1817 | N | ··· | ··· |
| CS70135 | E1818 | N | ··· | ··· |
| CS70136 | E1819 | N | ··· | ··· |
| CS70137 | E1825 | N | ··· | ··· |
| CS70138 | E1828 | N | ··· | ··· |
| CS70139 | E1832 | N | ··· | ··· |
| CS70140 | E1833 | N | ··· | ··· |
| CS70141 | E1853 | N | ··· | ··· |
| CS70142 | E1868 | N | ··· | ··· |
| CS70143 | E1950 | N | ··· | ··· |
| CS70144 | E1998 | N | ··· | ··· |
| CS70145 | E2034 | N | ··· | ··· |
| CS70146 | E217 | N | ··· | ··· |
| CS70147 | E562 | N | ··· | ··· |
| CS70148 | E1001 | N | ··· | ··· |
| CS70149 | E1368 | N | ··· | ··· |
| CS70150 | E1690 | N | ··· | ··· |
| CS70151 | E1704-1 | N | ··· | ··· |
| CS70152 | E1704-3 | N | ··· | ··· |
| CS70153 | E1715 | N | ··· | ··· |
| CS70154 | E1723 | N | ··· | ··· |
| CS70155 | E1735 | N | ··· | ··· |
| CS70156 | E1935 | N | ··· | ··· |
| CS70157 | E1967 | N | ··· | ··· |
| CS70158 | E2014 | N | ··· | ··· |
| CS70159 | E2057 | N | ··· | ··· |
| CS70160 | E2207 | N | ··· | ··· |
| CS70161 | E2406 | N | ··· | ··· |
| CS70162 | E2408 | Y | Y | Y |
| CS70163 | E2410 | N | ··· | ··· |
| CS70164 | E2415 | N | ··· | ··· |
| CS70165 | E2425 | N | ··· | ··· |
| CS70166 | E2425 | N | ··· | ··· |
| CS70167 | E2441 | N | ··· | ··· |
| CS70168 | E2443 | N | ··· | ··· |
| CS70169 | E2448 | N | ··· | ··· |
| CS70170 | E2491 | N | ··· | ··· |
| CS70171 | E2502 | N | ··· | ··· |
| CS70172 | E2513 | N | ··· | ··· |
| CS70173 | E2563 | N | ··· | ··· |
| CS70174 | E2609 | N | ··· | ··· |
| CS70175 | E2633 | N | ··· | ··· |
| CS70176 | E2676 | N | ··· | ··· |
| CS70177 | E2692 | Y | N | ··· |
| CS70178 | E2724 | N | ··· | ··· |
| CS70179 | E2763 | N | ··· | ··· |
| CS70180 | E2764 | N | ··· | ··· |
| CS70181 | E2779 | N | ··· | ··· |
| CS70182 | E2861 | N | ··· | ··· |
| CS70183 | E2862 | N | ··· | ··· |
| CS70184 | E2897 | N | ··· | ··· |
| CS70185 | E2904 | N | ··· | ··· |
| CS70186 | E2905 | N | ··· | ··· |
| CS70187 | E2947 | N | ··· | ··· |
| CS70188 | E2993 | N | ··· | ··· |
| CS70189 | E3004 | N | ··· | ··· |
| CS70190 | E3006 | N | ··· | ··· |
| CS70191 | E3017 | N | ··· | ··· |
| CS70192 | E3065 | N | ··· | ··· |
| CS70193 | E3134 | N | ··· | ··· |
| CS70194 | E3190 | N | ··· | ··· |
| CS70195 | E3198 | N | ··· | ··· |
| CS70196 | E3258 | N | ··· | ··· |
| CS70197 | E3267 | N | ··· | ··· |
| CS70198 | E3298 | N | ··· | ··· |
| CS70199 | E3313 | N | ··· | ··· |
| CS70200 | E3317 | Y | Y | N |
| CS70201 | E3430 | N | ··· | ··· |
| CS70202 | E3459 | N | ··· | ··· |
| CS70203 | E3462 | N | ··· | ··· |
| CS70204 | E3474 | N | ··· | ··· |
| CS70205 | E3478 | N | ··· | ··· |
| CS70206 | E3501 | N | ··· | ··· |
| CS70207 | E3505 | N | ··· | ··· |
| CS70208 | E3530 | N | ··· | ··· |
| CS70209 | E3531 | N | ··· | ··· |
| CS70210 | E3598-1 | N | ··· | ··· |
| CS70211 | E3598-2 | N | ··· | ··· |
| CS70212 | E3637 | N | ··· | ··· |
| CS70213 | E3642 | N | ··· | ··· |
| CS70214 | E3655 | Y | N | ··· |
| CS70215 | E3683 | N | ··· | ··· |
| CS70216 | E3700 | N | ··· | ··· |
| CS70217 | E3754 | N | ··· | ··· |
| CS70218 | E3756 | N | ··· | ··· |
| CS70219 | E3783 | Y | N | ··· |
| CS70220 | E3806 | N | ··· | ··· |
| CS70221 | E3816 | N | ··· | ··· |
| CS70222 | E3826 | N | ··· | ··· |
| CS70223 | E3876 | N | ··· | ··· |
| CS70224 | E3879 | N | ··· | ··· |
| CS70225 | E3880 | N | ··· | ··· |
| CS70226 | E3885 | Y | N | ··· |
| CS70227 | E3912 | Y | Y | Y |
| CS70228 | E3927 | N | ··· | ··· |
| CS70229 | E3930 | Y | N | ··· |
| CS70230 | E3963 | N | ··· | ··· |
| CS70231 | E3980 | N | ··· | ··· |
| CS70232 | E4009 | N | ··· | ··· |
| CS70233 | E4028 | Y | N | ··· |
| CS70234 | E4058 | N | ··· | ··· |
| CS70235 | E4096 | N | ··· | ··· |
| CS70236 | E4104 | N | ··· | ··· |
| CS70237 | E4105 | N | ··· | ··· |
| CS70238 | E4110 | N | ··· | ··· |
| CS70239 | E4118 | Y | N | ··· |
| CS70240 | E4129 | N | ··· | ··· |
| CS70241 | E4148 | N | ··· | ··· |
| CS70242 | E4150 | N | ··· | ··· |
| CS70243 | E4151 | N | ··· | ··· |
| CS70244 | E4162 | N | ··· | ··· |
| CS70245 | E4223 | N | ··· | ··· |
| CS70246 | E4247 | N | ··· | ··· |
| CS70247 | E4256 | N | ··· | ··· |
| CS70248 | E4272 | N | ··· | ··· |
| CS70249 | E4285 | N | ··· | ··· |
| CS70250 | E4295 | Y | Y | Y |
| CS70251 | E4350 | N | ··· | ··· |
| CS70252 | E4396 | N | ··· | ··· |
| CS70253 | E4411 | N | ··· | ··· |
| CS70254 | E4423 | N | ··· | ··· |
| CS70255 | E4491 | N | ··· | ··· |
| CS70256 | E4506 | Y | N | ··· |
| CS70257 | E4522 | Y | N | ··· |
| CS70258 | E4583 | N | ··· | ··· |
| CS70259 | E4589 | N | ··· | ··· |
| CS70260 | E4633 | N | ··· | ··· |
| CS70261 | E4680 | N | ··· | ··· |
| CS70262 | E4695 | N | ··· | ··· |
| CS70263 | E4715 | N | ··· | ··· |
| CS70264 | E4716 | Y | Y | Y |
| CS70265 | E4722 | Y | Y | Y |
| CS70266 | E4751 | N | ··· | ··· |
| CS70267 | E4791 | N | ··· | ··· |
| CS70268 | E4801 | N | ··· | ··· |
| CS70269 | E4811 | N | ··· | ··· |
| CS70270 | E4812 | N | ··· | ··· |
| CS70271 | E4820 | N | ··· | ··· |
| CS70272 | E4856 | Y | N | ··· |
| CS70273 | E4907 | N | ··· | ··· |
| CS70274 | E4930 | N | ··· | ··· |
| CS70275 | E4940 | N | ··· | ··· |
| CS70276 | E4970 | N | ··· | ··· |
| CS70277 | E5008 | N | ··· | ··· |
| CS70278 | E5025 | N | ··· | ··· |
| CS70279 | E5026 | N | ··· | ··· |
| CS70280 | E5085 | N | ··· | ··· |
| CS70281 | E5096 | Y | N | ··· |

N, no. Y, yes.
