## Supplemental Table 2 for "GAL4/GFP enhancer-trap lines for identification and manipulation of cells and tissues in developing Arabidopsis leaves"

**Table S2. Reproducibility of expression and pattern features**

| **Figure** | **Panel** | **No. leaves with displayed features / no. analyzed leaves** | **Assessed expression or pattern features** |
| --- | --- | --- | --- |
| 1 | K | 15/18 | Ubiquitous |
| 1 | L | 15/17 | Ubiquitous |
| 1 | M | 19/19 | Ubiquitous |
| 1 | N | 33/33 | Ubiquitous |
| 1 | O | 26/29 | Inner cells |
| 1 | P | 29/29 | Vascular cells in top half of primordium, inner cells in basal half of primordium |
| 1 | Q | 31/31 | Vascular cells in top half of primordium, inner cells in basal half of primordium |
| 1 | R | 19/19 | Vascular cells in top half of leaf, inner cells in basal half of leaf |
| 1 | S | 16/19 | Abaxial inner cells |
| 1 | T | 34/36 | Abaxial inner cells & middle tissue layer |
| 1 | U | 24/25 | Abaxial inner cells & middle tissue layer |
| 1 | V | 34/34 | Abaxial inner cells & middle tissue layer |
| 1 | X | 14/14 | Inner, nonvascular cells |
| 1 | Y | 14/14 | Inner, nonvascular cells |
| 1 | Z | 14/14 | Inner, nonvascular cells |
| 2 | A | 15 (adaxial) or 26 (abaxial) / 28 | Top third of adaxial epidermis & whole abaxial epidermis |
| 2 | B, left | 22/23 | Top three-quarters of epidermis & trichomes |
| 2 | B, right | 30/30 | Whole epidermis |
| 2 | C, left | 14/14 | Top three-quarters of epidermis & trichomes |
| 2 | C, right | 15/15 | Whole epidermis |
| 2 | D, left | 16/16 | Epidermis of whole lamina and petiole midline & trichomes |
| 2 | D, right | 18/18 | Whole epidermis |
| 2 | E | 16/16 | Trichomes |
| 2 | F | 17/18 | Top three-quarters of marginal epidermis |
| 2 | G | 14/14 | Whole marginal epidermis |
| 2 | H | 16/16 | Whole marginal epidermis |
| 2 | I | 59/59 | Whole epidermis |
| 2 | J, left | 42/42 | All cells of marginal epidermis, except few cells in top half of primordium |
| 2 | J, right | 45/45 | Whole epidermis |
| 2 | K, left | 33/38 | Bottom quarter and few cells in top three-quarters of marginal epidermis |
| 2 | K, right | 21/21 | Whole epidermis, including stomata |
| 2 | L, left | 31/31 | Bottom quarter and few cells in top three-quarters of marginal epidermis |
| 2 | L, right | 21/21 | Whole epidermis, including stomata |
| 2 | M | 29/30 | Absent |
| 2 | N | 26/26 | Top quarter of primordium |
| 2 | O | 18/18 | Top three-quarters of primordium |
| 2 | P | 18/18 | Whole leaf |
| 2 | Q | 31/33 | Absent |
| 2 | R | 19/21 | Top quarter of primordium |
| 2 | S | 23/28 | Top half of lamina |
| 2 | T | 16/18 | Top three-quarters of lamina |
| 3 | A | 22/22 | Midvein |
| 3 | B | 30/30 | Midvein |
| 3 | C | 16/17 | Midvein & first loop |
| 3 | D | 34/48 | Midvein & first and second loop |
| 3 | E | 25/25 | Absent |
| 3 | F | 20/20 | Midvein |
| 3 | G | 27/37 | Midvein & first loop |
| 3 | H | 24/28 | Midvein & first and second loop |
| 4 | A | ND | Narrow midvein & scalloped vein-network outline |
| 4 | B | 19/20 | Shapeless vascular cluster |
| 4 | C | 32/46 | Midvein & first and second loop |
| 4 | D | 21/21 | Shapeless vascular domain |
| 4 | E | 16/23 | Midvein & first and second loop |
| 4 | F | 18/18 | Broad vascular domain |
| 4 | G | 21/21 | Narrow midvein & scalloped vein-network outline |
| 4 | H | 19/19 | Broad vascular zone |

ND: not determined.
